# Inversion frequencies and phenotypic effects are modulated by the environment

**DOI:** 10.1101/295568

**Authors:** Emma Berdan, Hanna Rosenquist, Keith Larson, Maren Wellenreuther

**Affiliations:** Department of Marine Sciences, University of Gothenburg, Gothenburg, Sweden; MEMEG, Ecology Building, Lund University, Lund, Sweden; Department of Ecology and Environmental Sciences, Climate Impacts Research Centre, Umea University, Umeå, Sweden; The New Zealand Institute for Plant & Food Research Ltd, Auckland, New Zealand; School of Biological Sciences, the University of Auckland, New Zealand

**Keywords:** *Coelopa frigida*, inversion polymorphism, adaptation, GxE interactions, frequency effects

## Abstract

Understanding how environmental variation drives phenotypic diversification within species is a major objective in evolutionary biology. The seaweed fly *Coelopa frigida* provides an excellent model for the study of genetically driven phenotypes because it carries an α/β inversion polymorphism that affects body size. *Coelopa frigida* inhabits highly variable beds of decomposing seaweed on the coast in Scandinavia thus providing a suitable test ground to investigate the genetic effects of substrate on both the frequency of the inversion (directional selection) and on the phenotype (genotype x environment effects). Here we use a reciprocal transplant experiment to test the effect of the α/β inversion on body size traits and development time across four suitable natural breeding substrates from the clinal distribution. We show that while development time is unaffected by GxE effects, both the frequency of the inversion and the relative phenotypic effects of the inversion on body size differ between population x substrate combinations. This indicates that the environment modulates the fitness as well as the phenotypic effects of the inversion karyotypes. It further suggests that the inversion may have accumulated qualitatively different mutations in different populations that interact with the environment. Together our results are consistent with the idea that the inversion in *C. frigida* likely evolves via a combination of local mutation, GxE effects, and differential fitness of inversion karyotypes in heterogeneous environments.

## Introduction

One of the central goals of evolutionary biology is to understand how organisms cope with environmental heterogeneity (Savolainen et al. 2013). Phenotypic plasticity can solve this problem but the phenotypic response to variation in an environmental variable (i.e. phenotypic plasticity) may vary among species as well as among genotypes of the same species. This is commonly referred to as a Genotype-by-Environment interaction (GxE, Falconer 1990; Lynch and Walsh 1998; Via and Lande 1985). Abundant experimental data and theoretical work now provide strong support for the idea that the relative performance of different genotypes across environmental gradients may be involved in the maintenance of phenotypic plasticity and genetic variation as well as the evolution of fitness related phenotypes (Carreira et al. 2006; Ungerer et al. 2003).

The study of environmental heterogeneity has been championed by work on geographic patterns along clines, with classic examples coming from genetic clines in flies almost a century ago (Dobzhansky 1948; Sturtevant 1921). Other examples include the genetic underpinnings of traits involved in adaptation to high altitude (Storz 2002) and pigmentation (Hoekstra and Coyne 2007). Clines hold a large attraction as natural Darwinian laboratories because they allow researchers to quantify the gradual effects of changing environments on genotype, phenotype, and the interaction between them (Endler 1977). These studies have shown that while many environmental tolerance traits that vary clinally are quantitative and polygenic (Wellenreuther and Hansson 2016), they can also be located within chromosomal segments that segregate as a single locus (Tigano and Friesen 2016). If environmentally adaptive variants are chromosomally linked, either because they are found on chromosomal inversions or are in areas of low recombination, their selective coefficients act in an additive manner and they may be less likely to be lost via genetic drift and more likely to contribute to rapid adaptation (Kirkpatrick and Barton 2006). In the extreme case, complex adaptations may be accomplished through the fixation of linked sets of adaptive variants (Yeaman 2013), often called supergenes, a case in which polygenic adaptation may be more accurately modeled with single-locus models (Schwander et al. 2014). Good examples for such single-locus supergene adaptations come from *Drosophila melanogaster* where inversion polymorphisms produce latitudinally varying phenotypes for traits including heat and cold tolerance and body size (Reinhardt et al. 2014; Schrider et al. 2016).

Chromosomal inversion polymorphisms are being identified at a fast rate from an increasing number of populations of organisms, e.g. insects, plants, bacteria and humans, and are commonly linked to trait and fitness variation (Hoffmann and Rieseberg 2008; Kirkpatrick and Barton 2006). Inversions are particularly common in flies, and an intriguing mystery is why there are huge differences in levels of polymorphism and rates of fixation for inversions between closely related species and even between chromosomes in *Drosophila* spp. (Kirkpatrick 2010). The seaweed fly *Coelopa frigida* (family Coelopidae) and its sister species *C. nebularum* have a large chromosomal α/β inversion polymorphism on chromosome I encompassing about 10% of the genome and containing ∼100 genes (Aziz 1975). The inversion polymorphism is known to affect a wide range of life history and reproductive traits, such as development time (Day and Buckley 1980), body size (Butlin et al. 1982a), longevity and female fecundity (Butlin and Day 1984a; Butlin et al. 1982c), female resistance to male mating attempts (Gilburn and Day 1994; Gilburn et al. 1992) and also male’s ability to overcome female resistance (Butlin et al. 1982a).

The most striking of these phenotypic traits is a three-fold size difference in males: male homokaryotypes for the inversion type α are much larger (∼9mm) than β- homokaryotypes (∼3mm), whereas the heterokaryotype αβ is intermediate in size (∼6mm) (Figure 1a, Wellenreuther pers. obs., Butlin and Day 1984b). Female size is not significantly affected by karyotype (Butlin and Day 1984b). Maintenance of the polymorphism is secured by strong heterosis (Collins 1978), since αβ-heterokaryotypes have higher fitness than either homokaryotype (Butlin and Day 1984a; Butlin and Day 1989). The heterokaryotype αβ has a higher viability during development than either homokaryotype, both in experiments (Butlin and Day 1984a; Leggett et al. 1996) and in the field, as indicated by heterokaryotype excess (Butlin et al. 1982c; Day et al. 1983; Day et al. 1982b; Gilburn and Day 1994). Heterokaryotype excess increases under interspecific competition with the congener *C. pilipes* and under intense intraspecific competition (Butlin and Day 1984a; Leggett et al. 1996). The effect is disproportionally strong in males, especially decreasing the frequency of αα-homokaryotypes (Butlin and Day 1984a), possibly indicating lower viability of males of this karyotype. However, inversion frequencies also form a cline from the North Sea to the Baltic Sea, with the α inversion becoming more rare towards the Baltic Sea (Day et al. 1983).

**Figure 1:**
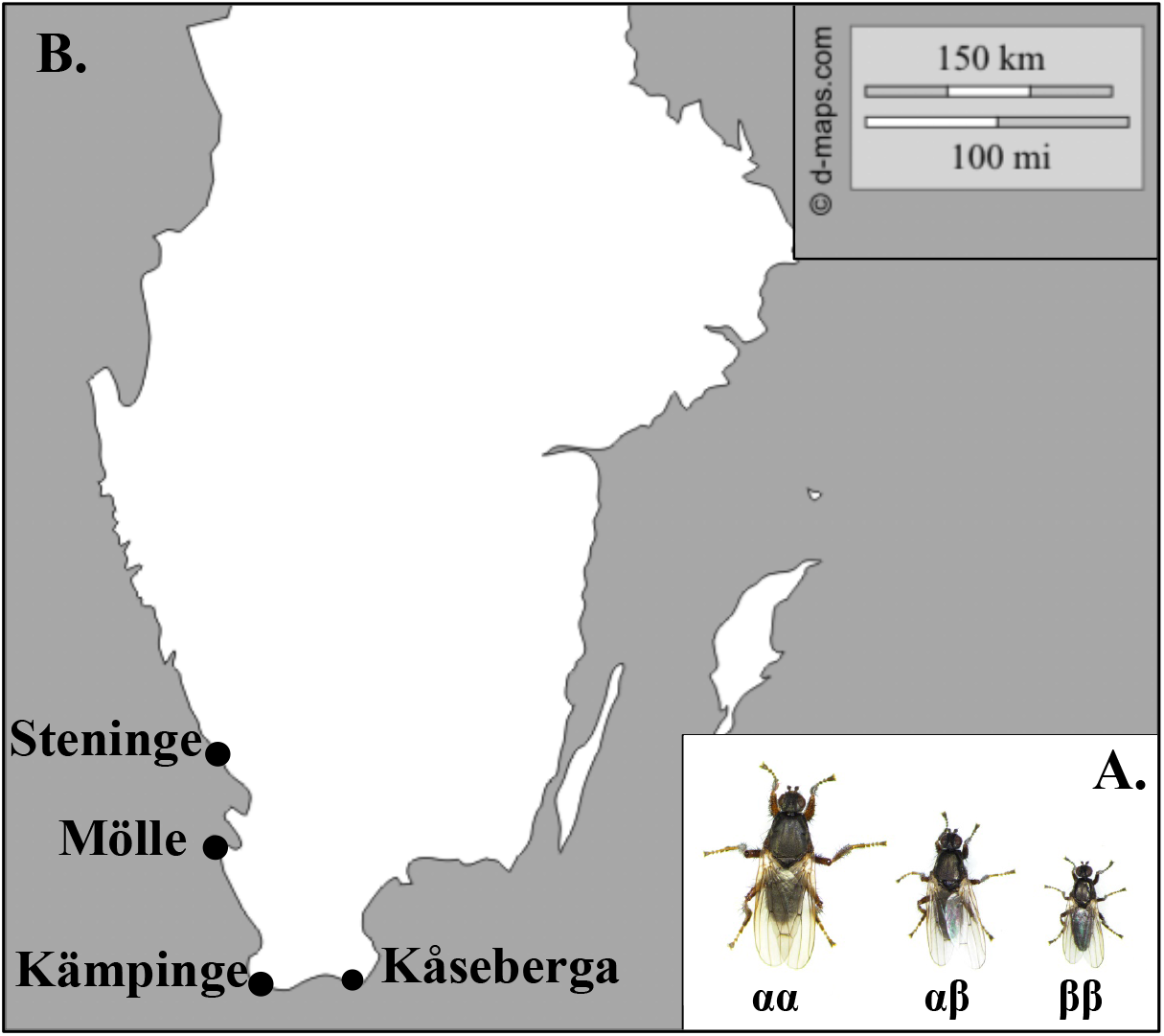
A. Picture of an αα, αβ, and ββ male showcasing the size differential. B. Map of *C. frigida* fly populations used in this study. Base map from d-maps (http://d-maps.com/carte.php7num_car=23102).

Here we explore if the environment modulates inversion frequencies and phenotypic effects in *C. frigida*. For this, we study *C. frigida* populations in Scandinavia that show variation in karyotype frequencies (Day et al. 1983). Specifically, we test the possible effects of the α/β inversion polymorphism on body traits and development time across four suitable natural breeding substrates using a fully balanced reciprocal transplant experiment. We also elucidate the relative fitness of the inversion karyotypes on these varying substrates by examining the frequencies of the three karyotypes in different environments.

## Materials and methods

### Study species and sites

*Coelopa frigida* belongs to the group of acalyptrate flies which exclusively forage on decomposing seaweed (Cullen et al. 1987) and is found all along the seashores of Scandinavia (Mcalpine 1991). This species plays a vital role in coastal environment health by accelerating the decomposition of algae, allowing for faster release of nutrients (Cullen et al. 1987), and are an important food source for predatory coastal invertebrates and seabirds (Fuller 2003; Summers et al. 1990). *Coelopa frigida* has 5 chromosomes with inversions occuring on 3 of them (Aziz 1975). The largest of these is the α/β which occurs on chromosome I. The α/β inversion occurs at stable frequencies in all populations so far studied in Northern Europe (Butlin et al. 1982a), although there may be seasonal (Butlin 1983) and clinal variation (Day et al. 1983).

Inversion frequencies of *C. frigida* form a cline from the North Sea to the Baltic Sea, with the α inversion becoming more rare towards the Baltic Sea (Day et al. 1983). Such clines are classic signs of selection and this cline is associated with a decrease in salinity and tidal range and associated changes in the composition of the wrackbed (i.e. the species of algae present). As salinity decreases from the North Sea to the Skagerrak and Kattegat regions there is a corresponding decrease of marine species of brown seaweeds such as *Fucus spp*. and *Laminaria spp*. moving into the Baltic Sea, *Zostera* eelgrass and red algae gradually displace the brown seaweeds (Table 1). We selected four populations (Kämpinge, Kåseberga, Mölle and Steninge) from the southern part of the cline that differ in tidal range, salinity, and wrackbed composition (Figure 1b, Table 1). We extracted climate data (sea surface salinity and air temperature) for each site from MARSPEC (Sbrocco and Barber 2013) and WorldClim databases using the R package raster (Hijmans and van Etten 2014) (Table 1).

**Table 1.** Spatial coordinates (WGS 1984 DD), sea surface salinity (PSU), mean air temperature, inversion frequency, and wrackbed composition of the study sites.

|  | Coordinates (WGS84, DD) |  | Sea surface salinity (PSU)<sup>a</sup> |  | Air Temperature (°C)<sup>b</sup> | Inversion frequency<sup>c</sup> | Wrackbed composition |  |  |  |
| --- | --- | --- | --- | --- | --- | --- | --- | --- | --- | --- |
| | Latitude | Longitude | Mean annual | Annual SD | Mean annual | Frequency of the $\alpha$ inversion | Percentage *Fucus* spp. | Percentage Kelp<sup>d</sup> | Percentage red algae<sup>e</sup> | Percentage eelgrass<sup>f</sup> |
| Steninge | 56.77 | 12.63 | 15.79 | 16.62 | 7.5 | 0.49 | 50 | 20 | 30 | 0 |
| Mölle | 56.28 | 12.49 | 15.38 | 16.88 | 7.8 | 0.42 | 30 | 30 | 30 | 10 |
| Kämpinge | 55.4 | 12.98 | 12.82 | 11.74 | 8.1 | 0.35 | 30 | 0 | 30 | 40 |
| Kåseberga | 55.38 | 14.06 | 9.91 | 5.4 | 8.0 | 0.34 | 30 | 0 | 50 | 20 |

<sup>a</sup> Data source: Sbrocco & Barber (2013), accessed on 1 November 2016.
<sup>b</sup> Data source: WorldClim database (1.4) , accessed in May 2018
<sup>c</sup> Day et al. (1983).
<sup>d</sup> *Saccharina* and *Laminaria* spp.
<sup>e</sup> *Furcellaria lumbricalis*
<sup>f</sup> *Zostera marina*

### Reciprocal transplant experiment

We conducted a fully balanced reciprocal transplant experiment involving all four locations, where larvae from all locations were allowed to grow on its own wrackbed substrate and the substrates of the remaining three locations. For each location, fly larvae and wrackbed substrate were collected between February and May 2014. The wrackbed substrate was placed in labelled plastic bags and then frozen to ensure that no viable eggs or larvae remained. Fly larvae from were reared on 150g of their own substrate in aerated plastic containers until a puparium was formed, and then kept in individual Eppendorf tubes (to ensure virginity) containing a small piece of cotton soaked in 5% sucrose solution. When a fly eclosed, the Eppendorf was moved to the refrigerator to slow down aging. *Coleopa frigida* is exceptionally cold tolerant and this treatment does not appear to affect survival (E. Berdan, pers. obs.). Over a maximum of ten days, adult flies were collected until a minimum of 25 couples could be set up per location. Eggs were collected from 14-15 mating pairs per population and subsequently mixed to populate three replicates of each population x substrate combination with 30 eggs each (4 populations x4 substrates x 3 replicates, each one with 30 eggs, total eggs for the experiment: 1440). Replicate pots were kept at 25° C with a 12/12 hour light:dark cycle and each pot (190×110×70 mm) was aerated and contained 150 g of seaweed. Pots were checked daily and all eclosing flies were sexed and then stored at −20° C on the same day.

### Morphological measurements

Morphological measurements were carried out on adult flies only. All flies were weighed (Sartorius Quintix 124-1S microbalance) to the nearest 0.0001 grams. The thorax width of each fly was measured with a digital calliper (Cocraft Digital Vemier Caliper) at the widest place and to the nearest 0.01 mm. This measurement was repeated twice and the average was used. Then, the left wing and right middle leg were removed (if one side was damaged the other side was used). The wing and leg were placed under a thin glass slide and placed under the microscope and a picture was the taken using a Nikon Digital Sight DS-Fi2 mounted on a Nikon SM2800 microscope set to a magnification of two power. The resulting image was labelled with an individual identifier. Measurements were conducted blind to location and substrate type to avoid bias.

### Molecular determination of inversion types

In *C. frigida* the two most common alcohol dehydrogenase (Adh) allozymes, Adh-B and Adh-D are in complete linkage disequilibrium with the α and β forms of the inversion (Day et al. 1982a). Gel electrophoresis was used to visualise the alcohol-dehydrogenase allozymes associated with the two inversion types and this allowed for straightforward genotyping of the inversion types. Starch gels (12%) were prepared by combining 24.5 g potato starch (Sigma-Aldrich, S4251) with 90 ml TEB-buffer and 90 ml dH_2_O, heating the mixture, and then pouring it onto an 18.5×10 cm glass plate with a plastic frame and left to set for 30 minutes, after which it was put in a 4°C refrigerator overnight.

Porcelain plates with 12 wells were kept on ice packs. On each well, a fly was placed together with a small amount of carborundum powder (Fisher Chemical, #C192-500) and one drop of cold dH_2_O. The flies were then homogenized using a glass rod. Once homogenized, filter paper wicks were placed in the wells to soak up the homogenate. The wicks were then gently inserted into the starch gel.

Starch gels were put in the electrophoresis chamber, which was connected to a power supply (Consort E863). The two chambers of the electrophoresis machine were filled with TEB-buffer. Two cloth wicks were soaked in TEB-buffer and placed so that they touched evenly along each of the exposed sides of the gel, while still connected to the buffer. An ice pack was placed on top of the gel and the electrophoresis (350 V, 50mA) took 1.5 hours.

After electrophoresis, the gel was cut horizontally with a metal thread and only the bottom half was used for staining. The agar overlay used for staining consisted of 20 ml of 2% agar, 10 ml HCl-Tris buffer (pH 8.6), 6 ml propan-2-ol, 1.5 ml 3-(4,5-dimethylthiazolyl-2)-2,5-diphenyltetrazolium bromide (MTT), 1 ml Phenazine methosulphate (PMS), and 10 ml Nicotinamide-adenine dinucleotide (NAD). The stain was mixed and poured onto the cut surface of the gel. The covered box with the gel was then carefully put in a dark heating cabinet at 37° C for 40 minutes, or until the banding pattern was clear and then photographed.

### Statistical analysis

We analyzed weight and development time (in days) in R (R Development Core Team 2015) using general linear models. Weight was highly correlated with all other morphological characters (Supplemental table S1) so we only analyzed weight data and development time in a statistical framework. Weight data was normally distributed and so was analyzed assuming a Gaussian distribution. Development time is essentially count data and right skewed, so we analyzed the data assuming a Poisson distribution. For both data sets we considered individual flies to be our unit of replication and used the lme4 package in R (Bates et al. 2014) to construct a model with all factors (population, substrate, sex, and karyotype) and all two and three way interactions. We also included pot (replicate) nested within population*substrate as a random factor. We ran the ‘Anova’ function in the R package car (Fox and Weisberg 2011) on the full models and then examined the F-values for each term. Starting with a model with only an intercept we built up a series of nested models adding a single main effect, two or three way interaction at a time by decreasing F-value (i.e. starting with the term with the highest F-value). We then used the ‘Anova’ function with the *X^2^* distribution to compare the models in a nested fashion (i.e. each model is only compared to the one above it) and used a *X^2^* test to determine if they were significantly different. We selected the model with the lowest AIC whose nested models were not significantly better.

Visual inspection of our morphological data indicated that males were more variable than females. We formally tested for differences in variability for each morphological measurement in a statistical framework using Krishnamoorthy and Lee’s (2014) modified signed-likelihood ratio test (MS-LRT) implemented in the R package cvequality (Marwick and Krishnamoorthy 2016). We used a Bonferroni correction to account for multiple testing (Bland and Altman 1995).

We further examined the relative proportions of the three karyotypes (αα, αβ, and ββ) in all population x substrate combinations. For these analyses we used pot as our unit of replication. We tested for differences in two ways. First, we tested for karyotype differences using a *X*^2^ test on a 3 karyotypes x 16 population-substrate combination matrix. Second, we used a Dirichlet regression model in the R package DirichletReg (Maier 2014). The Dirichlet regression model allows us to analyze karyotypes (variables with a bounded interval that sum up to a constant - in our case frequencies of karyotypes in a population) exhibiting skewness and heteroscedasticity, without having to transform the data (Maier 2014). We used a nested approach starting with a simple model and then adding factors. Models were tested against the one above them in a nested fashion using the ‘Anova’ function in the R package car (Fox and Weisberg 2011) to determine the best model.

## Results

### Phenotypic data

We had different sample sizes for different treatment combinations (see figure legends for Figures 3 and 4) due to differential survival of the three karyotype categories. Three of the 96 combinations had zero flies (ex: αα Kämpinge females in Kämpinge) but we do not expect that this unduly influenced our results as we did not investigate any 4 way interactions in any of our statistical models and thus did not examine these three combinations directly. All morphological characteristics were highly and significantly correlated with each other (Supplementary Table S1).

We only examined weight in a statistical framework, as this morphological trait is a good proxy of the other morphological measurements (Pearson correlation coefficients with other traits varied from 0.78-0.89, Supplemental Table S1) while still being an easily tractable and understandable measurement. Weight was not strongly correlated with development time (Pearson’s product-moment correlation: 0.10) and therefore development time was examined separately. F-values from the full models indicated that effect sizes were much larger for weight than development time (Table S2). The best model for weight included all main effects, the two-way interactions: sex*karyotype, population*sex, population*karyotype, population*substrate, substrate*karyotype, and the three-way interaction: location*sex*karyotype. Comparing the best model with the next most simplified model gave a *X*^2^ value of 43.3 (*P* = 0.00073). The best model for development time included the main effects and the two-way interaction karyotype*sex. Comparing the best model with the next most simplified model gave a *X*^2^ value of 8.6 (*P* = 0.0135). The best models for both weight and development time included all main effects and a subset of interactions (Table 2). However, a GxE interaction (as evidenced by a population x substrate interaction) was only present for morphological characteristics, not for development time.

**Table 2.** Best models for adult weight and development time. An X indicates that a main effect or interaction was present in the best model.

|  | Main Effects |  |  |  | 2-way interactions |  |  |  |  | 3-way interactions |  |
| --- | --- | --- | --- | --- | --- | --- | --- | --- | --- | --- | --- |
|  | Karyotype | Population | Sex | Substrate | Karyotype:Population | Karyotype:Sex | Karyotype:Substrate | Population:Sex | Population:Substrate | Sex:Substrate | Karyotype:Population:Substrate |
| Weight | x | x | x | x | x | x | x | x | x |  | x |
| Development Time | x | x | x | x |  | x |  |  |  |  |  |

For all measured traits, the Krishnamoorthy and Lee’s (2014) modified signed-likelihood ratio test (MS-LRT) showed that males had significantly more phenotypic variation than females (Table 3). This variation could be partially accounted for by karyotype. Previous work has shown that males are on average bigger than females and that within male size differences are primarily driven by the αβ inversion system: αα males are the biggest followed by αβ and ββ (Butlin et al. 1982c). This held true in our male data set (Figure 2) but the magnitude of these differences depended on both population and substrate type (Figure 2, Supplementary Figures 2-5). Although the relative sizes and size differences of the karyotypes differed between combinations of location and substrate, the pattern of both leg length and wing length, thorax width, and weight (Figure 2, Supplementary Figures 2-5) in relation to karyotype and sex was clear and consistent for males, but much less pronounced for females. Likewise, across treatments (combination of population and substrate), αα-males had the largest mean size, followed by αβ-males and ββ-males in all different population x substrate combinations (Figure 2). For females, less than half of the combinations of population and substrate followed the same pattern.

**Table 3.**
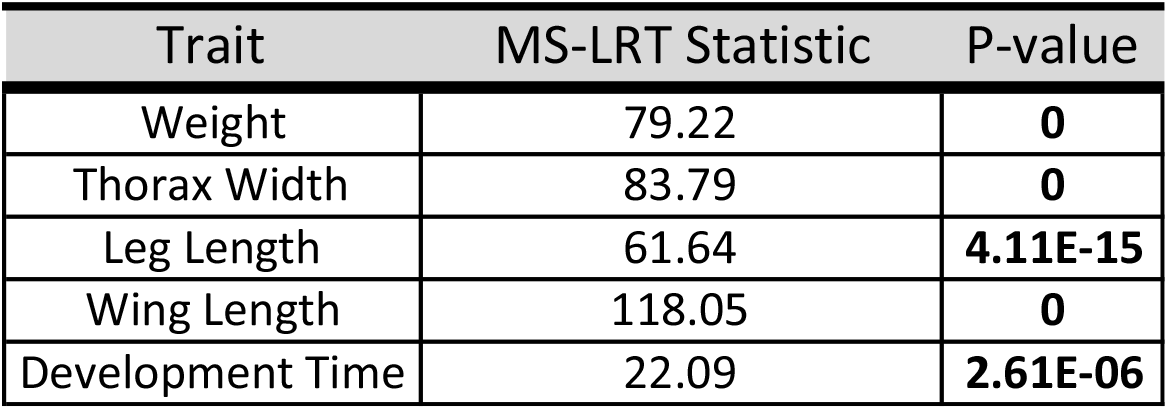
Results from Krishnamoorthy and Lee’s (2014) modified signed-likelihood ratio test (MS-LRT) looking to see if the coefficient of variation is different between males and females. Bolded p-values are significant at the P<0.0001 level after a Bonferroni correction.

| Trait | MS-LRT Statistic | P-value |
| --- | --- | --- |
| Weight | 79.22 | **0** |
| Thorax Width | 83.79 | **0** |
| Leg Length | 61.64 | **4.11E-15** |
| Wing Length | 118.05 | **0** |
| Development Time | 22.09 | **2.61E-06** |

**Figure 2:**
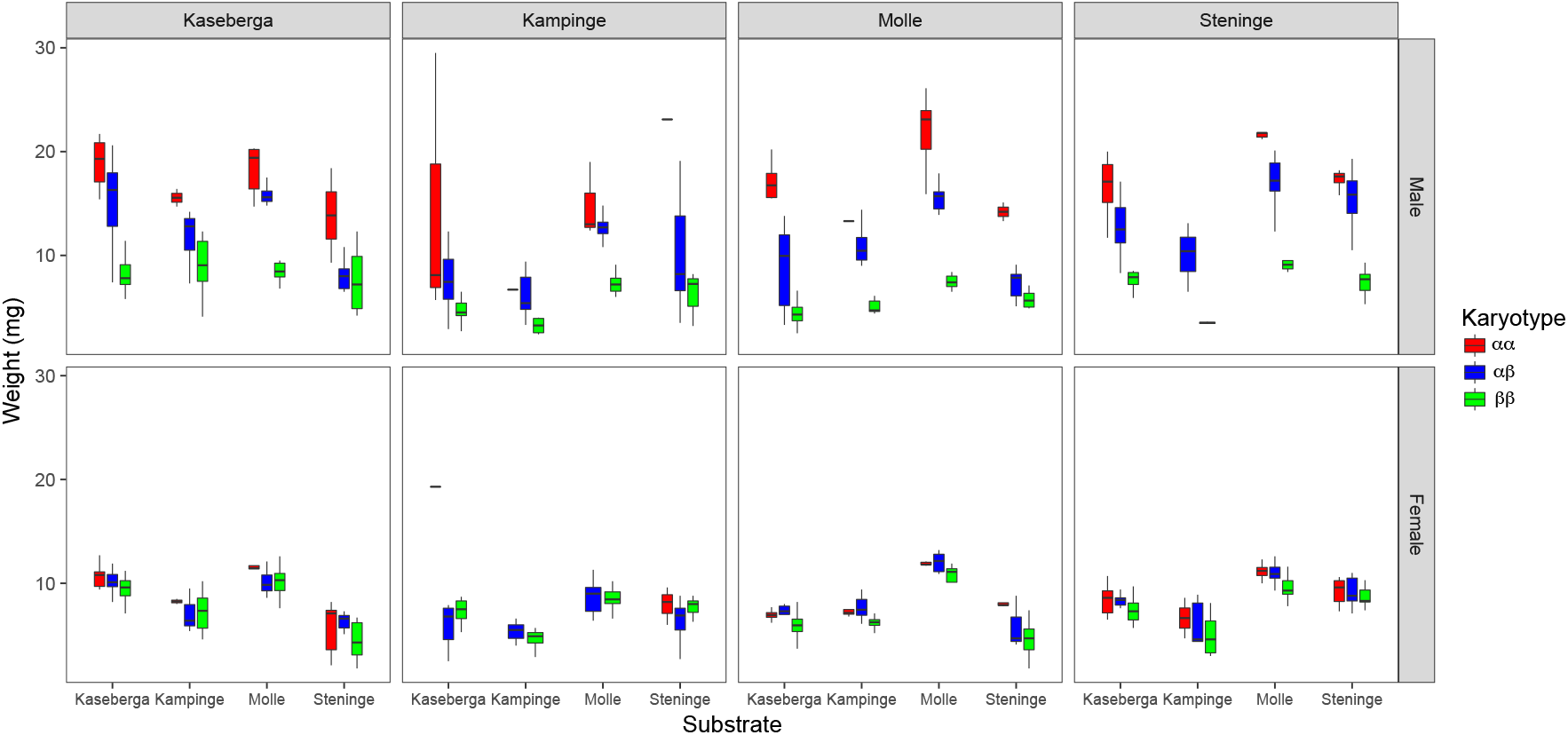
Boxplot of weight across populations (listed at Top), substrate (listed on x-axis), sex (indicated by top (male) and bottom (female) charts), and karyotype (indicated by color, red - αα, blue - αβ, and green-ββ). Outliers are not shown for clarity. N for each population x sex combination (from left to right for each karyotype and substrate as listed in the figure): Kåseberga Male: 10, 19, 5, 2, 11, 6, 5, 9, 10, 2, 9, 6, Kämpinge Male: 3, 20, 15, 1, 9, 4, 3, 13, 19, 1, 21, 6, Mölle Male: 10, 14, 1, 6, 8, 13, 6, 12, 13, 2, 6, 10, Steninge Male: 7, 17, 7, 0, 3, 2, 5, 15, 5, 7, 16, 14, Kåseberga Female: 7, 24, 11, 2, 3, 6, 4, 12, 18, 5, 13, 5, Kämpinge Female: 1, 12, 11, 0, 5, 4, 0, 13, 18, 3, 17, 9, Mölle Female: 4, 9, 4, 4, 10, 4, 6, 6, 5, 4, 3, 8, Steninge Female: 8, 21, 8, 2, 5, 4, 8, 19, 6, 7, 21, 7.

Development time did not follow the same pattern as observed for morphology (Figure 3). While αα-males generally took longer to develop, this was not always the case for every combination of population and substrate. For example, ββ flies from Steninge, raised on Kämpinge substrate took longer to develop than any other flies from Steninge. For females, the MS-LRT analysis demonstrated that there was considerably less variation between karyotypes compared to males and patterns varied substantially between populations and substrates.

**Figure 3:**
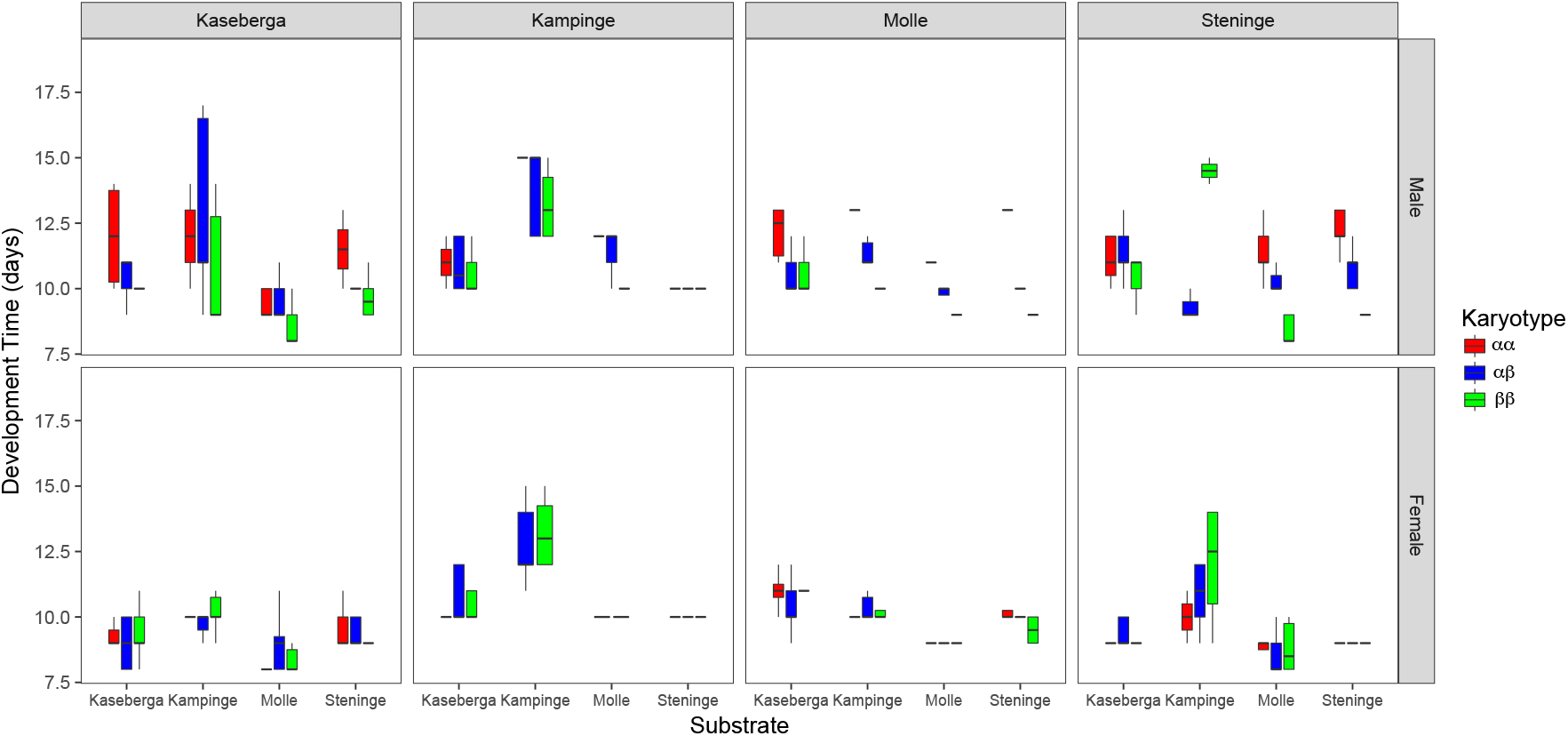
Boxplot of development time across populations (listed at Top), substrate (listed on x-axis), sex (indicated by top (male) and bottom (female) charts), and karyotype (indicated by color, black - αα, grey - αβ, and white-ββ). Outliers are not shown for clarity. N for each population x sex combination (from left to right for each karyotype and substrate as listed in the figure): Kåseberga Male: 10, 19, 5, 2, 11, 6, 5, 9, 10, 2, 9, 6, Kämpinge Male: 3, 20, 15, 1, 9, 4, 3, 13, 19, 1, 21, 6, Mölle Male: 10, 14, 13, 1, 6, 8, 6, 12, 13, 2, 6, 10, Steninge Male: 7, 17, 7, 0, 3, 2, 5, 15, 5, 7, 16, 14, Kåseberga Female: 7, 24, 11, 2, 3, 6, 4, 12, 18, 5, 13, 5, Kämpinge Female: 1, 12, 11, 0, 5, 4, 0, 13, 18, 3, 17, 9, Mölle Female: 4, 9, 4, 4, 10, 4, 6, 6, 5, 4, 3, 8, Steninge Female: 8, 21, 8, 2, 5, 4, 8, 19, 6, 7, 21, 7.

### Inversion types

The frequencies of αα, αβ, and ββ were significantly different across the Population x Substrate combinations (*X*^2^ = 71.5, df=30, *P* < 0.0001). Generally αβ was the most frequently observed karyotype, but ββ could be just as frequent or more frequent depending on the substrate. In ¾ of the populations (all except Steninge) ββ was the most frequent karyotype in at least one substrate and that was usually the substrate from Mölle.

We used Dirichlet regression to see which parameters best predicted the karyotype frequencies. With this model we were not able to examine F-values so we compared all possible models with the ‘Anova’ function from the R package car (Fox and Weisberg 2011). The best model was population + substrate + population*substrate. Comparing this with the next most simplified model gave a *X*^2^ value of 73.99 (P = 0.0002).

## Discussion

Our results demonstrate that the environment modulates both the frequencies of the karyotypes as well as their phenotypic effects in *C. frigida*. The presence of a significant GxE term for both of these indicates that there is population specific variation in larval fitness of the different inversion types as well as in the phenotypic effects of the inversion. The same pool of eggs was used to seed every substrate treatment per population so the neutral expectation (without sampling bias) would be similar karyotype frequencies across substrates.

Instead, we found strong differences in karyotype frequencies between substrates with some trends being consistent across multiple populations (ex: higher frequency of ββ in Mölle substrate, Figure 4). While it is possible that these results are due simply to sampling bias, it is unlikely that sampling bias would produce consistent trends. Furthermore, this result confirms previous research showing that the substrate composition has a strong selective effect on the inversion (Butlin et al. 1982b; Day et al. 1983; Edward and Gilburn 2013; Mérot et al. in press). Karyotypic specific survival differed across populations suggesting that that population specific variation has accumulated in the inversion. For example, ββ was the most common karyotype on Mölle wrack for every population except Steninge (Figure 4). Instead, in Steninge, little variance was seen between substrate combinations compared to other populations.

**Figure 4:**
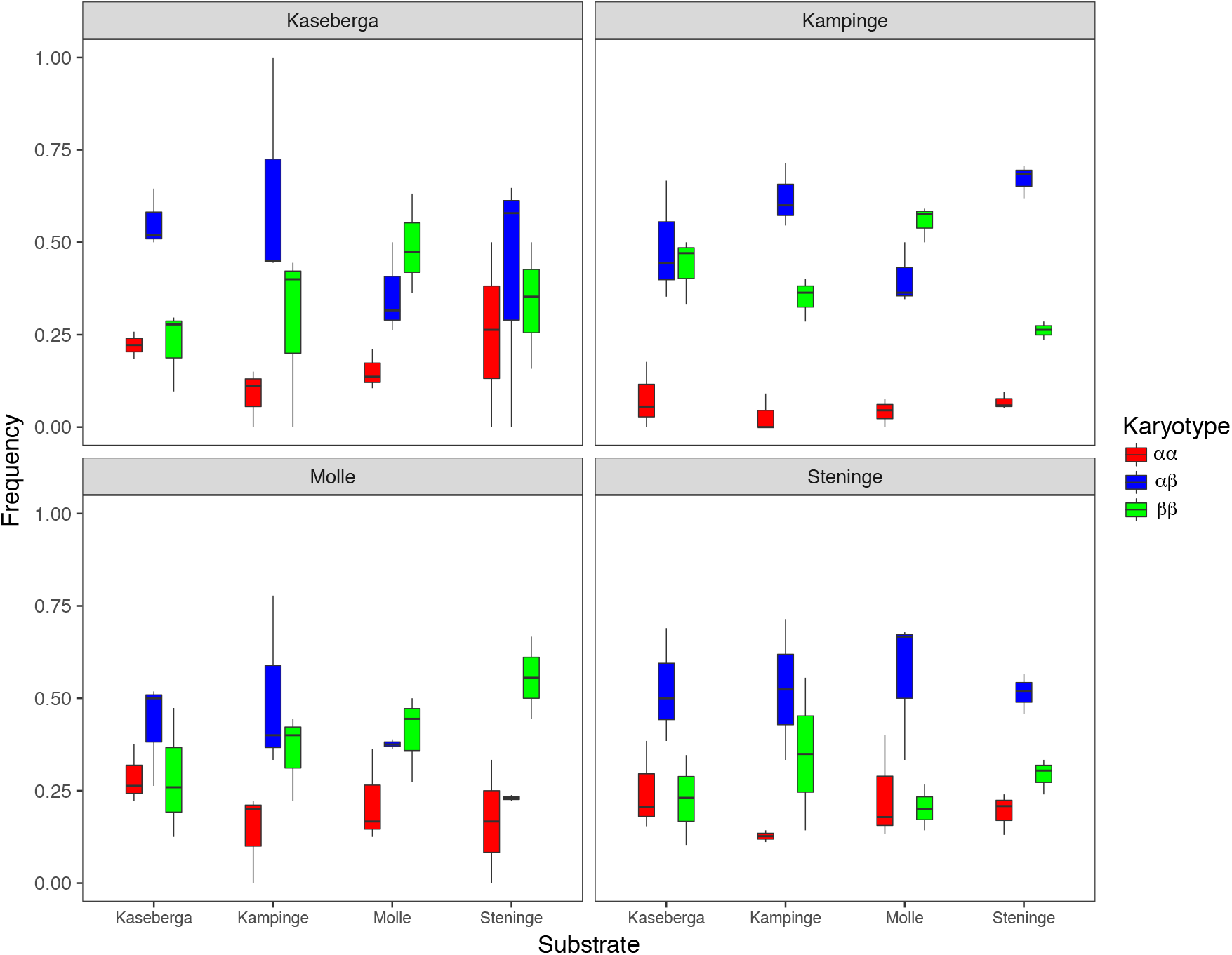
Box plot of frequencies of different karyotypes (black - αα, grey - αβ, and white-ββ) across populations (listed at the top of each graph) and substrates (listed on the x-axis).

Our results also indicate that inversion-related size effects in *C. frigida* are affected by the environment, particularly in males as females show little to no effect of karyotype on size. Male fly inversion phenotypes from the same population could vary two-fold for weight when grown on the same substrate (e.g. Steninge population on Mölle substrate), and three-fold when comparing male inversion phenotypes across all four growing substrates (e.g. Steninge, Figure 2). Male size has a high heritability with virtually all the variance in size being attributable to the chromosomal inversion system (Wilcockson et al. 1995). In all of our treatment combinations, environmental effects never swamped the effect of the inversion. Male αα were always the biggest, followed by αβ and ββ, which is consistent with previous work (Figure 1a; e.g. Butlin et al. 1982a). Population and substrate changed the magnitude of these differences as well as size averages but never the pattern of the inversion size effects on males. Part of this variation may be due to parental effects as the parents were reared in their “home” substrate. Parental larval diet has been shown to influence offspring size in *Drosophila melanogaster* (ex: Valtonen et al. 2012; Vijendravarma et al. 2010). However, despite strong environment and inversion effects on male size, the inversion, the population, and the environment exhibited negligible size effects in females. Specifically, our data show much lower between inversion variation in size and considerably less variation across environments for females. Thus, if maternal effects are present they are not influencing males and females in a similar manner. A similar difference in effect size between males and females has also been found in a common garden study on *C. frigida* grown on artificially mixed substrates containing in excess either the seaweed genus *Laminaria* or *Fucus* (Edward and Gilburn 2013). The study by Edward and Gilburn (2013) also found that flies grown on an excess of *Laminaria* showed greater among inversion size differences compared to *Fucus*, suggesting that the type of seaweed itself, or the associated microbiome of the seaweed, causes phenotypic differences in size. The wrackbed composition differs markedly along the populations studied, as a consequence of the declining salinity from the North Sea to the Baltic Sea (See Figure 1 and 2 in Wellenreuther et al. 2017), and have likely been major selective environmental factors in modulating *C. frigida* populations. Development time, another trait known to be affected by inversion type, also showed significantly higher variation across male karyotypes, but exhibited less extreme environmentally induced effects compared to size, and consequently no significant GxE effects. These results together with previous studies (Butlin and Day 1984a; Day et al. 1980) underscore that the inversion related phenotype effects in *C. frigida* are significantly affected by sex, with male phenotypes being more labile compared to female phenotypes.

Interactions between genotypes and the environment are commonly observed in quantitative traits (Gupta and Lewontin 1982), particularly those associated with fitness (e.g. Fanara et al. 2006), and are thought to provide a potent force for maintaining genetic variation (Fernández Iriarte and Hasson 2000; Gillespie and Turelli 1989; Hedrick 1986). The habitat used by *C. frigida* is heterogeneous and the resulting local population structure is similar to Levene’s migration model of subdivided populations, namely by a single pool of individuals that mate at random and breed on discrete and ephemeral resources. Environmental fluctuations in seaweed substrate composition likely favors the persistence of alternative rearrangements (i.e. the α/β inversion) containing sets of co-adapted alleles via balancing selection (Carroll et al. 2001; Filchak et al. 2000; Weinig 2000), with each set being optimal under different conditions (Dobzhansky 1970). This hypothesis may account for the so called “supergene” idea (Schwander et al. 2014) such as those determining mating types found in heterostylous plants and wing mimicry polymorphisms found in *Heliconius* butterflies (Thompson and Jiggins 2014). Balancing selection resulting from seasonal and stochastic temporal and spatial wrackbed changes may thus be a significant contributor to maintaining the fitness related inversion variation in size. Other examples for selection on fitness trade-offs come from introduced apple *Malus pumila* and native hawthorn *Crataegus* spp. hosts which has led to divergence in phenology and thermal physiology in the true fruit fly *Rhagoletis pomonella* (Filchak et al. 2000).

Balancing selection has also been implicated in the maintenance of other inversion polymorphisms, but whether balancing selection can maintain genetic polymorphisms over long time scales is still debated (Bernatchez 2016). Theoretical studies indicate that balancing selection may primarily be a short term process because the genetic load that it creates then has a negative feedback that hinders the long-term maintenance of adaptive alleles (Fijarczyk and Babik 2015). Furthermore, rapid environmental changes may cause dramatic allele frequency shifts that could cause the loss of formerly stable polymorphisms (Fijarczyk and Babik 2015). Despite this, recent studies have provided some evidence that balancing selection can operate even over longer time scales. Lindtke et al. (2017) investigated the inversion related cryptic colour polymorphism in *Timema cristinae* walking stick insects using a comprehensive dataset of natural populations and population genetic analyses. They were able to demonstrate that inversion colour variation has likely been maintained over extended time periods by balancing selection, which has previously been thought to be only possible over short time frames. Balancing selection (via spatially variable selection) on chromsomal inversions has been widely demonstrated in *Drosophila* species (Wellenreuther and Bernatchez 2018). The most well studied of these is the ln(3R)Payne inversion in *Drosophila melanogaster*, which shows stable parallel clines pertaining to latitude on 3 continents (Kapun et al. 2016). In *Drosophila pseudoobscura* many different arrangements on the third chromosome form clines across the Southwestern United States that are maintained by a migration selection balance between different environmental niches (Schaeffer 2008).

In conclusion, our results indicate that the inversion in *C. frigida* likely evolves via a combination of local mutation, GxE effects, and differential fitness of karyotypes in heterogeneous environments. Future work should apply high-resolution genome-wide approaches that target the regions inside the inversion to identify the true genic targets of varying selection. This approach would also elucidate which genes inside the inversion are linked to these large size effects in males, and how the environment has shaped the differentiation between these genes along gradients.

## Acknowledgements

MW would like to thank Roger Butlin and Andre Gilburn for helpful advice regarding the molecular inversion determination and rearing techniques. EB would like to thank Fabian Roger for helpful advice regarding the figures. Helena Westerdahl and Jan Åke Nilsson gave some helpful comments on earlier versions of this work. The work was supported by a grant from the Crafoord Foundation, from Entomological Foundation in Lund and from the Swedish Research Vetenskapsrådet Council (2012-3996) to MW. EB was supported by a Marie Skłodowska-Curie fellowship 704920 — ADAPTIVE INVERSIONS — H2020-MSCA-IF-2015.

## Conflict of Interest

The authors declare that they have no conflict of interest.

## Data Archiving

All raw phenotypic data and karyotype frequencies will be deposited on Dryad upon acceptance.

## Supplemental Material

**Table S1.** Pearson correlations of morphological factors.

|  | **Adult Weight** | **Thorax Width** | **Wing Length** | **Leg Length** |
| --- | --- | --- | --- | --- |
| **Weight** | 1 | 0.8306678 | 0.8850752 | 0.7786367 |
| **Thorax Width** | 0.8306678 | 1 | 0.8452453 | 0.7628986 |
| **Wing length** | 0.8850752 | 0.8452453 | 1 | 0.865498 |
| **Leg Length** | 0.7786367 | 0.7628986 | 0.865498 | 1 |

**Table S2.**
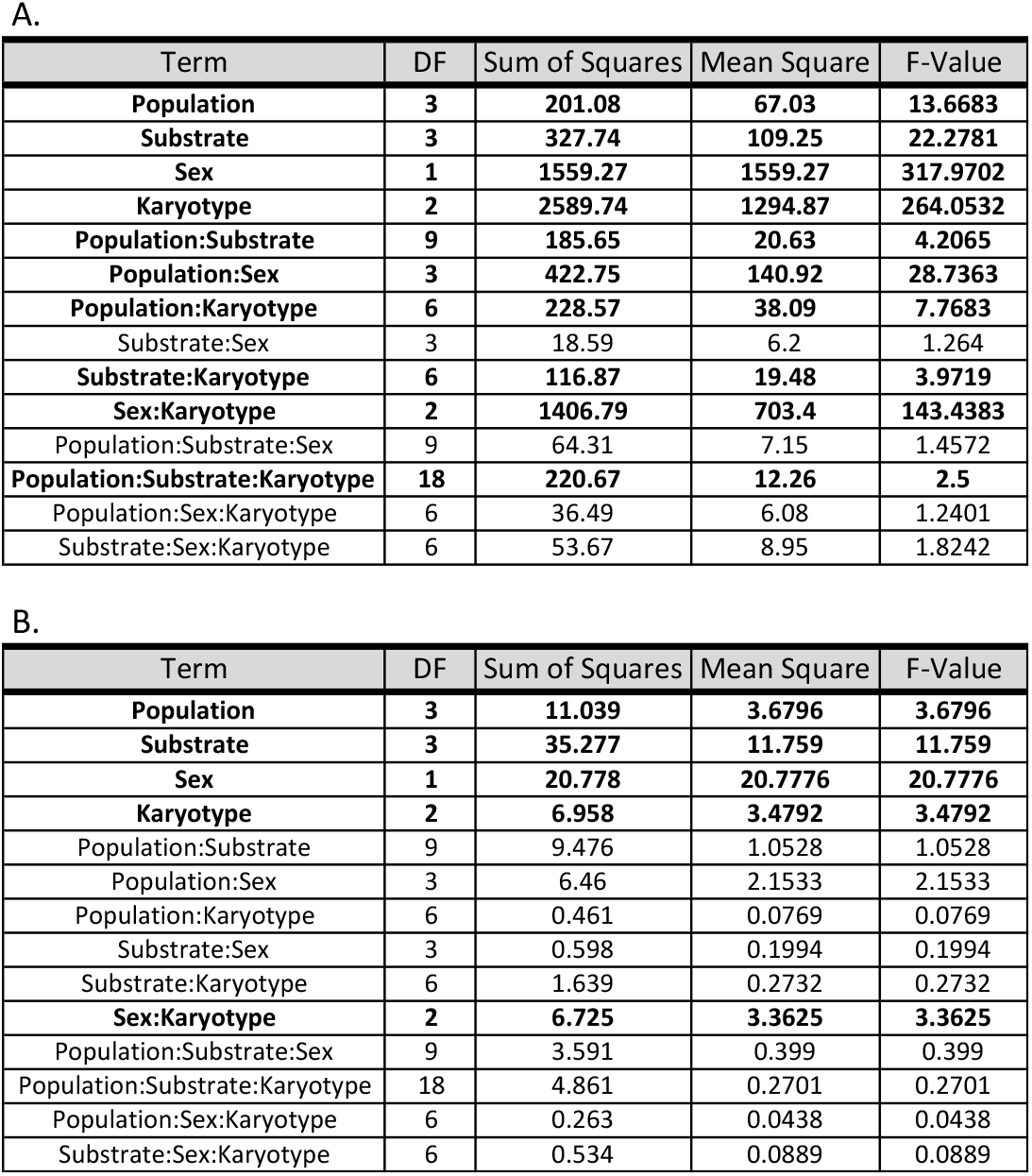
ANOVA tables for the full models adult weight (A) and development time (B). Bolded terms are included in the final models.

**Figure S2:**
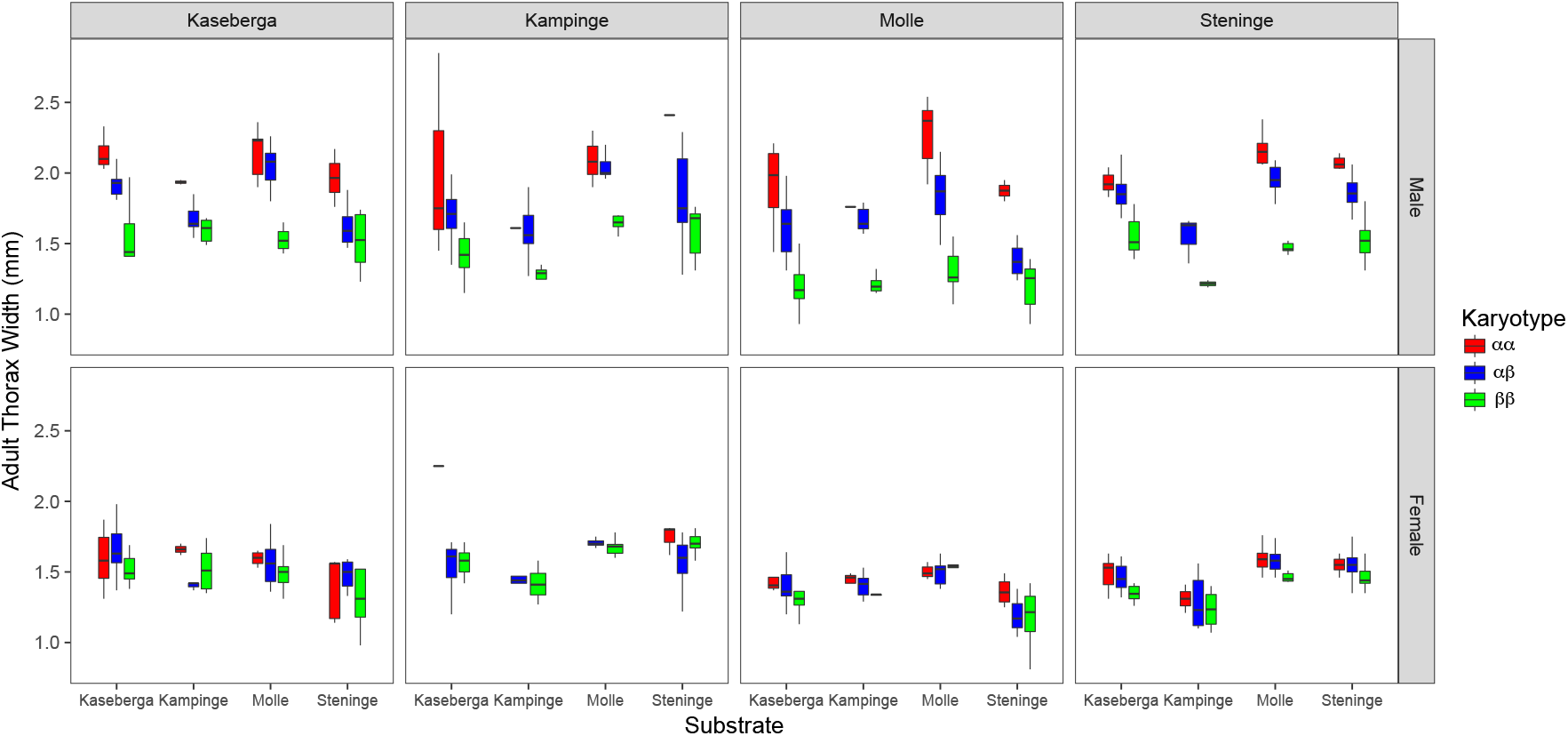
Boxplot of thorax width across populations (listed at Top), substrate (listed on x-axis), sex (indicated by top (male) and bottom (female) charts), and karyotype (indicated by color, red - αα, blue - αβ, and green-ββ). Outliers are not shown for clarity.

**Figure S3:**
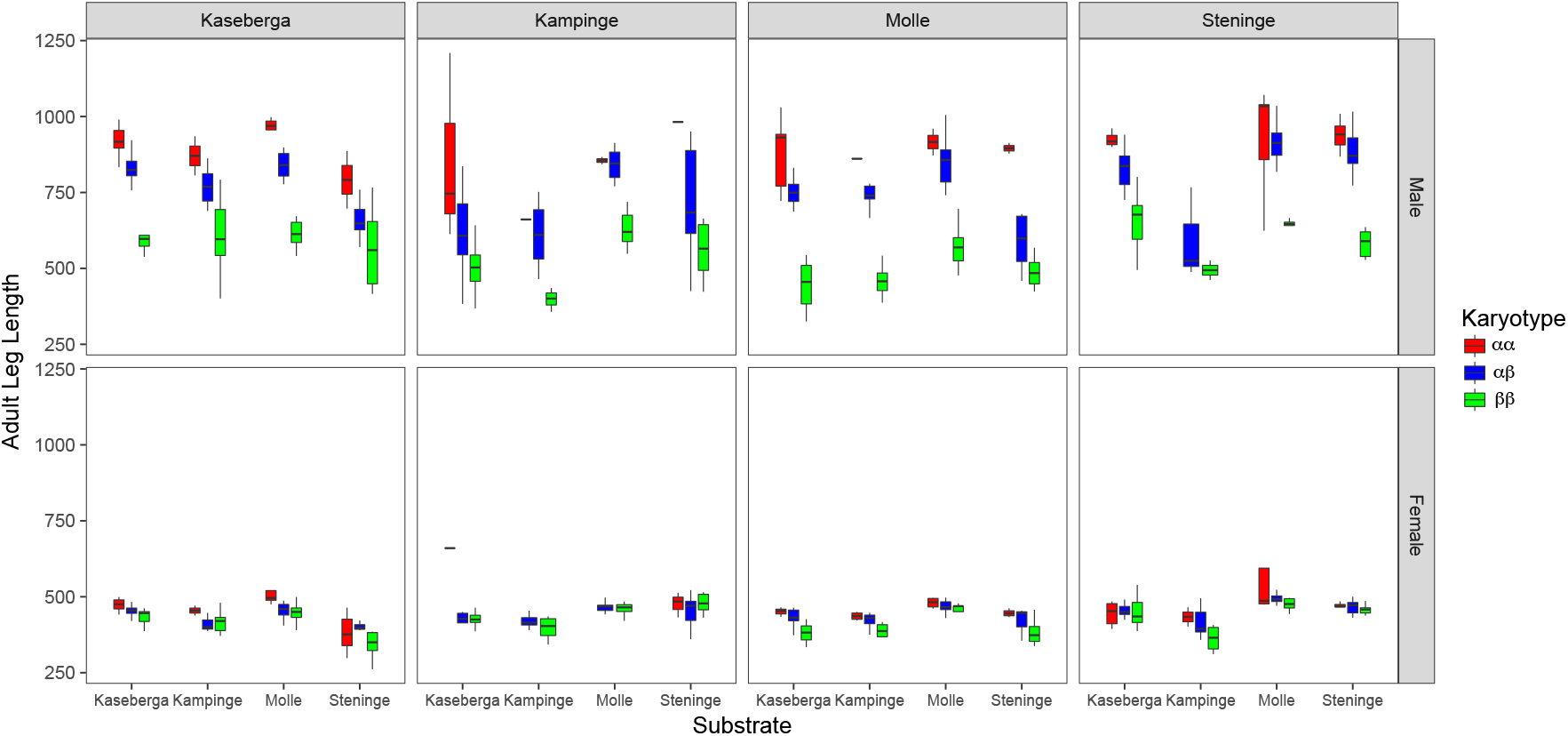
Boxplot of leg length across populations (listed at Top), substrate (listed on x-axis), sex (indicated by top (male) and bottom (female) charts), and karyotype (indicated by color, red - αα, blue - αβ, and green-ββ). Outliers are not shown for clarity.

**Figure S4:**
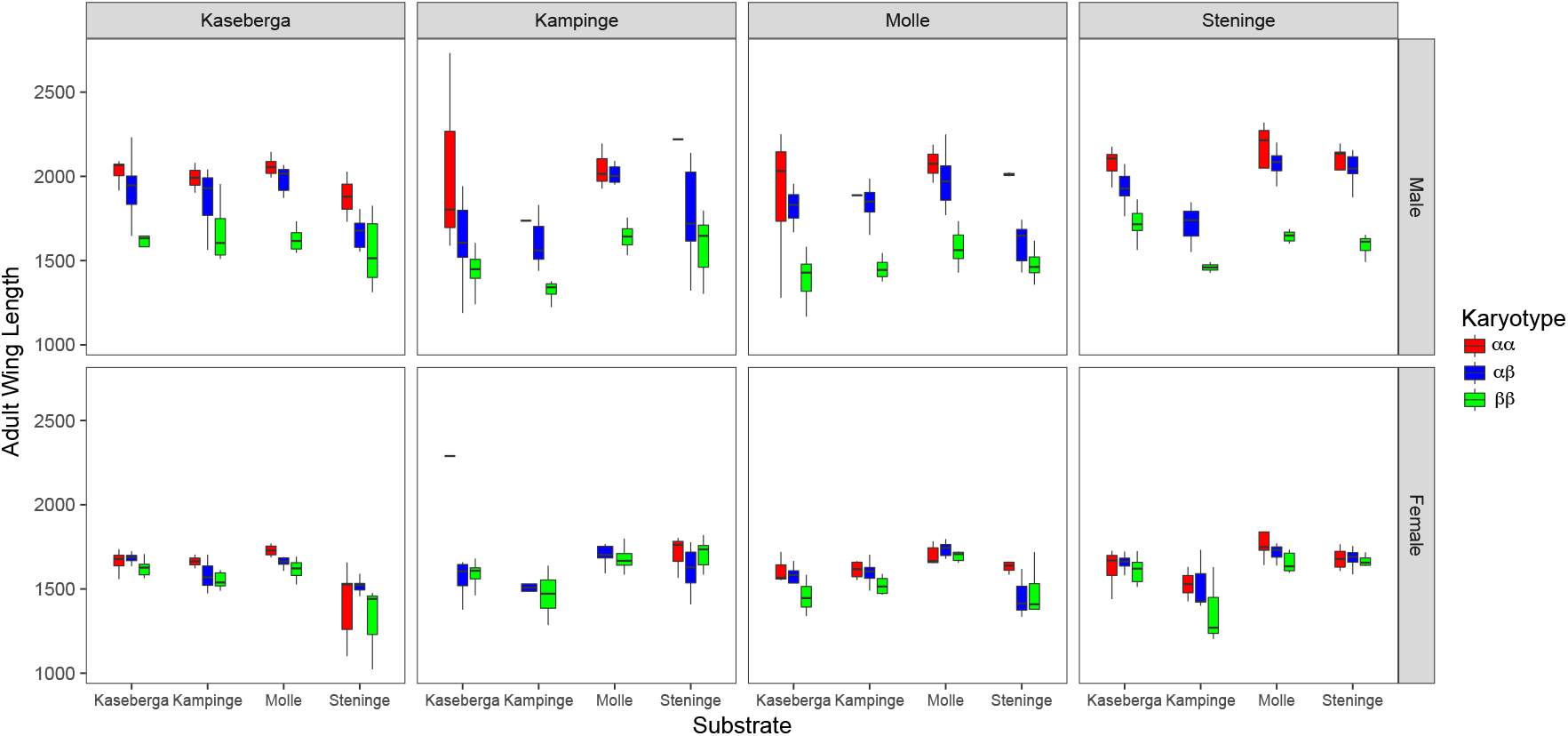
Boxplot of wing length across populations (listed at Top), substrate (listed on x-axis), sex (indicated by top (male) and bottom (female) charts), and karyotype (indicated by color, red - αα, blue - αβ, and green-ββ). Outliers are not shown for clarity.

## References

Aziz JB (1975) Investigations into chromosomes 1, 2 and 3 of Coelopa frigid (Fab.). Thesis,

Bates D, Mächler M, Bolker B, Walker S (2014) Fitting linear mixed-effects models using lme4 arXiv preprint arXiv:14065823

Bernatchez L (2016) On the maintenance of genetic variation and adaptation to environmental change: considerations from population genomics in fishes Journal of Fish Biology 89:2519–2556

Bland JM, Altman DG (1995) Multiple significance tests: the Bonferroni method Bmj 310:170

Butlin RK (1983) The maintenance of an inversion polymorphism in Coelopa frigida. University of Nottingham

Butlin RK, Collins P, Skevington S, Day T (1982a) Genetic variation at the alcohol dehydrogenase locus in natural populations of the seaweed fly, Coelopa frigida Heredity 48:45–55

Butlin RK, Collins PM, Skevington SJ, Day TH (1982b) Genetic-Variation at the Alcohol-Dehydrogenase Locus in Natural-Populations of the Seaweed Fly, Coelopa-Frigida Heredity 48:45–55 doi:DOI 10.1038/hdy.1982.5

Butlin RK, Day TH (1984a) The effect of larval competition on development time and adult size in the seaweed fly, Coelopa frigida Oecologia 63:122–127

Butlin RK, Day TH (1984b) The Effect of Larval Competition on Development Time and Adult Size in the Seaweed Fly, Coelopa-Frigida Oecologia 63:122–127 doi:Doi 10.1007/Bf00379793

Butlin RK, Day TH (1989) Environmental correlates of inversion frequencies in natural populations of seaweed flies (Coelopa frigida) Heredity 62:223–232

Butlin RK, Read IL, Day TH (1982c) The effects of a chromosomal inversion on adult size and male mating success in the seaweed fly, Coelopa frigida Heredity 49:51–62

Carreira VP, Soto IM, Hasson E, Fanara JJ (2006) Patterns of variation in wing morphology in the cactophilic Drosophila buzzatii and its sibling D. koepferae J Evol Biol 19:1275–1282 doi:10.1111/j.1420-9101.2005.01078.x

Carroll SP, Dingle H, Famula TR, Fox CW (2001) Genetic architecture of adaptive differentiation in evolving host races of the soapberry bug, Jadera haematoloma Genetica 112:257–272

Collins PM (1978) Studies on genetic polymorphism in *Coelopa frigida*. The Genetical Society of Great Britain,

Cullen SJ, Young AM, Day TH (1987) Dietary requirements of seaweed flies (Coelopa frigida) Estuarine, Coastal and Shelf Science 24:701–710

Day T, Dobson T, Hillier P, Parkin D, Clarke B (1982a) Associations of enzymic and chromosomal polymorphisms in the seaweed fly, Coelopa frigida Heredity 48:35–44

Day TH, Buckley PA (1980) Alcohol dehydrogenase polymorphism in the seaweed fly, Coelopa frigida Biochem Genet 18:727–742 doi:10.1007/bf00484589

Day TH, Dawe C, Dobson T, Hillier PC (1983) A chromosomal inversion polymorphism in Scandinavian populations of the seaweed fly, Coelopa frigida Hereditas 99:135–145 doi:10.1111/j.1601-5223.1983.tb00738.x

Day TH, Dobson T, Hillier PC, Parkin DT, Clarke B (1982b) Associations of enzymic and chromosomal polymorphisms in the seaweed fly, Coelopa frigida Heredity 48:35–44

Day TH, Dobson T, Hillier PC, Parkin DTa, Clarke B (1980) Different rates of development associated with the alcohol dehydrogenase locus in the seaweed fly, Coelopa frigida Heredity 44:321–326

Dobzhansky T (1948) Genetics of natural populations. XVI. Altitudinal and seasonal changes produced by natural selection in certain populations of *Drosophila pseudoobscura* and *Drosophila persimilis* Genetics 33:158

Dobzhansky TG (1970) Genetics of the evolutionary process. Columbia University Press, New York

Edward DA, Gilburn AS (2013) Male-specific genotype by environment interactions influence viability selection acting on a sexually selected inversion system in the seaweed fly, Coelopa frigida Evolution 67:295–302

Endler JA (1977) Geographic variation, speciation, and clines. Princeton University Press,

Falconer D (1990) Selection in different environments: effects on environmental sensitivity (reaction norm) and on mean performance Genetics Research 56:57–70

Fanara J, Folguera G, Iriarte PF, Mensch J, Hasson E (2006) Genotype by environment interactions in viability and developmental time in populations of cactophilic Drosophila Journal of Evolutionary Biology 19:900–908

Fernández Iriarte P, Hasson E (2000) The role of the use of different host plants in the maintenance of the inversion polymorphism in the cactophilic Drosophila buzzatii Evolution 54:1295–1302

Fijarczyk A, Babik W (2015) Detecting balancing selection in genomes: limits and prospects Molecular Ecology 24:3529–3545

Filchak KE, Roethele JB, Feder JL (2000) Natural selection and sympatric divergence in the apple maggot Rhagoletis pomonella Nature 407:739–742

Fox J, Weisberg S (2011) An R Companion to Applied Regression. 2 edn. SAGE Publications, Thousand Oaks, California

Fuller RA (2003) Factors influencing foraging decisions in ruddy turnstones Arenaria interpres (L.). Durham University

Gilburn AS, Day TH (1994) Sexual dimorphism, sexual selection and the α β chromosomal inversion polymorphism in the seaweed fly, Coelopa frigida Proceeding of the Royal Society of Biological Sciences 257:303–309

Gilburn AS, Foster SP, Day TH (1992) Female mating preference for large size in *Coelopa frigida* (seaweed fly) Heredity 69:209–216

Gillespie JH, Turelli M (1989) Genotype-environment interactions and the maintenance of polygenic variation Genetics 121:129–138

Gupta AP, Lewontin R (1982) A study of reaction norms in natural populations of Drosophila pseudoobscura Evolution 36:934–948

Hedrick PW (1986) Genetic polymorphism in heterogeneous environments: a decade later Annual review of ecology and systematics 17:535–566

Hijmans RJ, van Etten J (2014) raster: Geographic data analysis and modeling R package version 2

Hoekstra HE, Coyne JA (2007) The locus of evolution: evo devo and the genetics of adaptation Evolution 61:995–1016 doi:10.1111/j.1558-5646.2007.00105.x

Hoffmann AA, Rieseberg LH (2008) Revisiting the impact of inversions in evolution: from population genetic markers to drivers of adaptive shifts and speciation? Annual Review of Ecology, Evolution, and Systematics 39:21–42 doi:doi:10.1146/annurev.ecolsys.39.110707.173532

Kapun M, Fabian DK, Goudet J, Flatt T (2016) Genomic Evidence for Adaptive Inversion Clines in Drosophila melanogaster Mol Biol Evol 33:1317–1336 doi:10.1093/molbev/msw016

Kirkpatrick M (2010) How and why chromosome inversions evolve PLoS Biol 8:e1000501 doi:10.1371/journal.pbio.1000501

Kirkpatrick M, Barton N (2006) Chromosome inversions, local adaptation and speciation Genetics 173:419–434

Krishnamoorthy K, Lee M (2014) Improved tests for the equality of normal coefficients of variation Computational Statistics 29:215–232

Leggett M, Wilcockson R, Day T, Phillips D, Arthur W (1996) The genetic effects of competition in seaweed flies Biol J Linn Soc 57:1–11

Lindtke D et al. (2017) Long-term balancing selection on chromosomal variants associated with crypsis in a stick insect Molecular Ecology 26:xxx–xxx doi:10.1111/mec.14280

Lynch M, Walsh B (1998) Genetics and analysis of quantitative traits vol 1. Sinauer Sunderland, MA,

Maier MJ (2014) DirichletReg: Dirichlet regression for compositional data in R Research Report Series / Department of Statistics and Mathematics WU Vienna University of Economics and Business:125

Marwick B, Krishnamoorthy K (2016) cvequality: Tests for the Equality of Coefficients of Variation from Multiple Groups., R package version 0.1.1. edn.,

Mcalpine DK (1991) Review of the Australian Kelp Flies (Diptera, Coelopidae) Syst Entomol 16:29–84 doi:DOI 10.1111/j.1365-3113.1991.tb00573.x

Mérot C, Berdan EL, Babin C, Normandeau E, Wellenreuther M,Bernatchez L (in press) Inter-continental karyotype-environment parallelism supports a role for a chromosomal inversion in local adaptation in a seaweed fly Proccedings of the Royal Society B

R Development Core Team R (2015) R: A language and environment for statistical computing vol 1. R Foundation for Statistical Computing. R Foundation for Statistical Computing. doi:10.1007/978-3-540-74686-7

Reinhardt JA, Kolaczkowski B, Jones CD, Begun DJ, Kern AD (2014) Parallel geographic variation in Drosophila melanogaster Genetics 197:361–373

Savolainen O, Lascoux M, Merila J (2013) Ecological genomics of local adaptation Nature Review Genetics 14:807–820 doi:10.1038/nrg3522

Sbrocco EJ, Barber PH (2013) MARSPEC: ocean climate layers for marine spatial ecology Ecology 94:979–979

Schaeffer SW (2008) Selection in Heterogeneous Environments Maintains the Gene Arrangement Polymorphism of Drosophila Pseudoobscura Evolution 62:3082–3099 doi:10.1111/j.1558-5646.2008.00504.x

Schrider DR, Hahn MW, Begun DJ (2016) Parallel evolution of copy-number variation across continents in Drosophila melanogaster Molecular biology and evolution 33:1308–1316

Schwander T, Libbrecht R, Keller L (2014) Supergenes and complex phenotypes Current Biology 24:288–294 doi:10.1016/j.cub.2014.01.056

Storz JF (2002) Contrasting patterns of divergence in quantitative traits and neutral DNA markers: analysis of clinal variation Mol Ecol 11:2537–2551

Sturtevant AH (1921) A case of rearrangement of genes in *Drosophila* Proceedings of The National Academy of Science of the united States of America 7:235–237

Summers RW, Smith S, Nicoll M, Atkinson NK (1990) Tidal and Sexual Differences in the Diet of Purple Sandpipers Calidris-Maritima in Scotland Bird Study 37:187–194

Thompson M, Jiggins C (2014) Supergenes and their role in evolution Heredity 113:1–8

Tigano A, Friesen VL (2016) Genomics of local adaptation with gene flow Molecular ecology

Ungerer MC, Halldorsdottir SS, Purugganan MD, Mackay TF (2003) Genotype-environment interactions at quantitative trait loci affecting inflorescence development in Arabidopsis thaliana Genetics 165:353–365

Valtonen TM, Kangassalo K, Polkki M, Rantala MJ (2012) Transgenerational Effects of Parental Larval Diet on Offspring Development Time, Adult Body Size and Pathogen Resistance in Drosophila melanogaster Plos One 7 doi:ARTN e31611 10.1371/journal.pone.0031611

Via S, Lande S (1985) Genotype–environment interaction and the evolution of phenotypic plasticity Evolution 39:505–522

Vijendravarma RK, Narasimha S, Kawecki TJ (2010) Effects of parental larval diet on egg size and offspring traits in Drosophila Biol Lett 6:238–241 doi:10.1098/rsbl.2009.0754

Weinig C (2000) Plasticity versus canalization: population differences in the timing of shade-avoidance responses Evolution 54:441–451

Wellenreuther M, Bernatchez L (2018) Eco-Evolutionary Genomics of Chromosomal Inversions Trends Ecol Evol 33:427–440 doi:10.1016/j.tree.2018.04.002

Wellenreuther M, Hansson B (2016) Detecting polygenic evolution: problems, pitfalls, and promises Trends in Genetics 32:155–164 doi:10.1016/j.tig.2015.12.004

Wellenreuther M, Rosenquist H, Jaksons P, Larson W (2017) Local adaptation along an environmental cline in a species with an inversion polymorphism J Evol Biol 30:1068–1077

Wilcockson R, Crean C, Day T (1995) Heritability of a Sexually Selected Character Expressed in Both Sexes Nature 374:158–159

Yeaman S (2013) Genomic rearrangements and the evolution of clusters of locally adaptive loci Proceedings of the National Academy of Sciences doi:10.1073/pnas.1219381110

